# Chronic sleep curtailment expediates brain aging by activating the complement and coagulation cascades in mice

**DOI:** 10.1101/2025.02.18.638886

**Authors:** Pawan K. Jha, Utham K. Valekunja, Akhilesh B. Reddy

## Abstract

Chronic sleep insufficiency is prevalent in modern society and has been associated with age-related neurodegenerative diseases. Loss of sleep accelerates the progression of neurodegeneration in animal models of neurological diseases. Here, we study whether chronic sleep curtailment leads to brain aging in wild-type animals without a genetic predisposition. We used a wild-type mouse model to simulate modern-day conditions of restricted sleep and compared the brain (cortex) proteome of young sleep-restricted animals with different aged control groups. We report the alteration of 149 proteins related to sleep and 1269 related to age with 96 proteins common between them. Through pathway analysis of proteins common to sleep restriction and aging, we discovered that the complement and coagulation cascade pathways were enriched by alterations of complement component 3 (C3), alpha-2-macroglobulin (A2M), fibrinogen alfa and beta chain (FGA and FGB). This is the first study indicating the possible role of the complement and coagulation pathways in brain aging and by chronic sleep restriction (CSR) in mice.

## INTRODUCTION

The quality and average duration of sleep has decreased in modern society. Insufficient sleep is associated with long-term health consequences that include physiological and cognitive ailments, and this may lead to a shorter life expectancy ^1^. Sleep disruption exacerbates age-related neurodegenerative pathologies such as dementia and Alzheimer’s disease ^2^, and evidence emerging from epidemiological studies highlights that sleep disturbance is often present years before the symptomatic stage of neurodegenerative disease and worsens during its progression ^3,4^. Interestingly, Harrison and Rothwell showed that sleep-deprived young adults exhibit similar prefrontal neuropsychological dysfunction as alert people aged ∼ 60 years ^5^. Together, these important pieces of evidence point to the hypothesis that sleep loss accelerates aging or aging-like phenotypes. This hypothesis is supported by findings suggesting that sleep facilitates metabolite clearance, whereas aging reduces waste clearance within the brain ^6,7^.

A general decline in sleep efficiency and cognitive function are hallmarks of physiological aging ^8^. It is well established that age-associated changes are caused mainly by the loss of neuronal cells, reduced neuronal networks, and shortening of dendrites ^9–11^. Proteomic analyses of aging brain tissue from both humans and mice reveal coordinated molecular changes across key cellular pathways. These alterations manifest in proteins governing mitochondrial function, synaptic transmission, neuroinflammation, DNA repair, myelination, and apoptosis – creating a distinct molecular signature of neural aging ^12,13^. Chronic sleep restriction (CSR) has emerged as a critical factor in cognitive decline, particularly through its detrimental effects on memory function. Prior work revealed that CSR triggers brain inflammation and synapse deterioration ^14,15^ – neurological disruptions that strikingly parallel the hallmarks of aging. This connection between sleep insufficiency and brain aging has gained further support from recent neuroimaging evidence, which demonstrates that inadequate sleep accelerates age-related changes in brain structure and function ^16^. Despite these compelling observations linking sleep deprivation to accelerated brain aging, the underlying molecular mechanisms driving this relationship remain largely unexplored.

To investigate the molecular mechanisms underlying the emerging link between sleep deprivation and accelerated brain aging, we conducted a comprehensive proteomic analysis comparing chronic sleep restriction (CSR) and age-related changes in the mouse brain. Our study examined the cerebral cortex proteome of young mice subjected to sleep restriction, middle-aged mice with normal sleep patterns, and aged mice with normal sleep patterns. Through comparative analysis of differentially abundant proteins, we uncovered distinct molecular signatures. CSR predominantly altered the expression of proteins involved in cytoplasmic translation, vesicle-mediated transport, and metallothionein binding. In contrast, aging specifically affected proteins associated with neuronal projections, synaptic organization, mitochondrial fatty acid β-oxidation, and neutrophil degranulation. Remarkably, our pathway analyses revealed that both CSR and aging converged on common molecular pathways: the complement and coagulation cascades. This novel finding represents the first proteomic evidence highlighting the significance of these pathways in both CSR and aging-related brain changes. These shared molecular signatures provide a promising foundation for understanding how chronic sleep debt may accelerate brain aging, opening new avenues for therapeutic intervention.

## RESULTS

### Experimental design and proteome coverage

We investigated proteome changes using young C57BL/6J mice (3-4 months old) subjected to different sleep protocols, alongside age-related controls. Our sleep restriction paradigm consisted of five experimental groups: sleep-deprived young mice (SD) underwent forced locomotion for 20 hours (ZT4-ZT20) with 4 hours of sleep permitted for 5 days; recovery sleep mice (RS) were similarly sleep-deprived allowed them to recover for 2 days, on third day they experienced 10 hours of forced locomotion during their active period (ZT14-24) with 14 hours of sleep allowed; like normal sleep controls (NS). We also included two age-related control groups: middle-aged (MID) and old-aged (OLD) mice with normal sleep patterns (Figure 1A).

**FIGURE 1.**
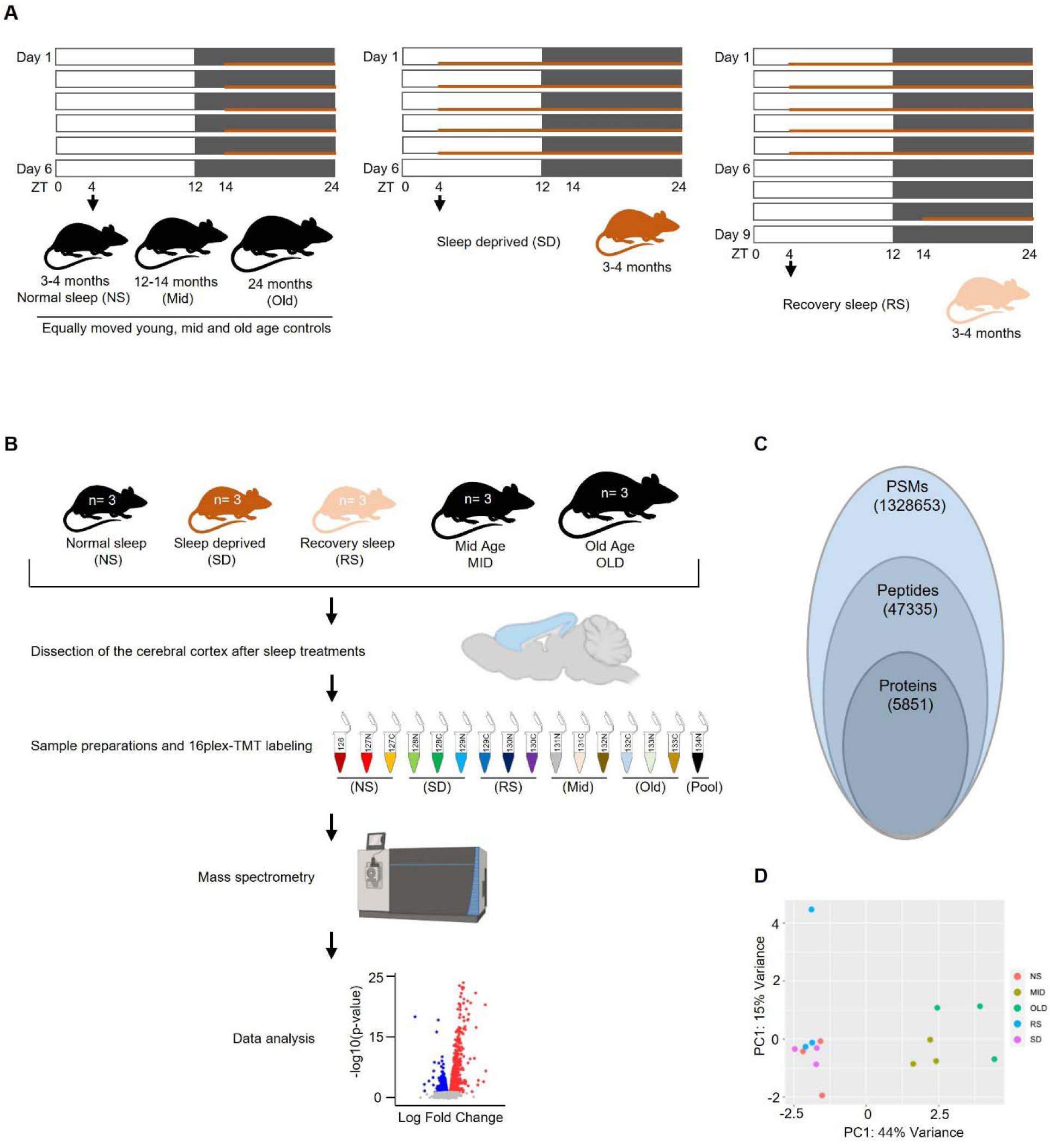
Schematic representation of experimental strategy, proteomics workflow, and quality control (QC). **(A)** Animals from SD group were subjected to sleep restriction for 20 h (ZT4-ZT24) by FA for 5 d (Day 1-Day 5) (*middle*), same age NS, MID, and OLD age control groups (*left*) were subjected to FA for 10h (ZT10-ZT14) at double speed and RS group was subjected to sleep restriction like SD group and allowed to recover for two days (Day 6-Day 7) and on Day 8 subjected to FA like controls (*right*). White and black bars represent light and dark periods and orange lines represent FA timelines. Arrow at ZT4 indicates termination of experiment followed by sacrifice and isolation of cerebral cortex. (**B**) Workflow for 16plex-TMT quantitative proteomics from cortex, from microdissection to bioinformatics analysis. Black color mice, sizes small to large: NS, MID, and OLD. Orange and light orange color: SD and RS group. (**C**) Overview of proteome coverage obtained in TMT-based quantitative analysis. (**D**) PCA plot showing segregations of all five groups, color-coded. NS: normal Sleep, SD: sleep deprived, RS: recovery sleep, MID: mid-age, OLD: old age, FA: forced ambulation.

To analyze proteomic changes, we harvested the cerebral cortex at ZT4, four hours after the final treatment period. We employed quantitative proteomics using 16-plex tandem mass tag (TMT) labeling coupled with liquid chromatography-tandem mass spectrometry (LC-MS/MS) (Figure 1B). This approach allowed simultaneous analysis of all experimental groups within a single mass spectrometry run, eliminating inter-run variability. With this approach, we identified 132,865 peptide spectral matches (PSMs) and 47,335 unique peptides, corresponding to 5,851 distinct proteins (Figure 1C). Principal component analysis (PCA) revealed clear segregation among all five experimental groups (NS, SD, RS, MID, and OLD), indicating distinct proteomic signatures for each condition (Figure 1D).

### CSR contributes to proteomic alteration in the cortex of young mice

We investigated how chronic sleep restriction (CSR) shapes protein dynamics in the cerebral cortex by examining protein expression in the young mice group, which were exposed to varying sleep conditions (Figure 2A-C). Our proteomic analysis revealed that CSR significantly modifies 149 proteins (2.5% of 5,737 proteins; q < 0.2). Notably, we found that CSR exerts more subtle effects on cortical protein expression than acute sleep deprivation, aligning with previous findings on sleep-dependent gene regulation ^17–19^.

**FIGURE 2.**
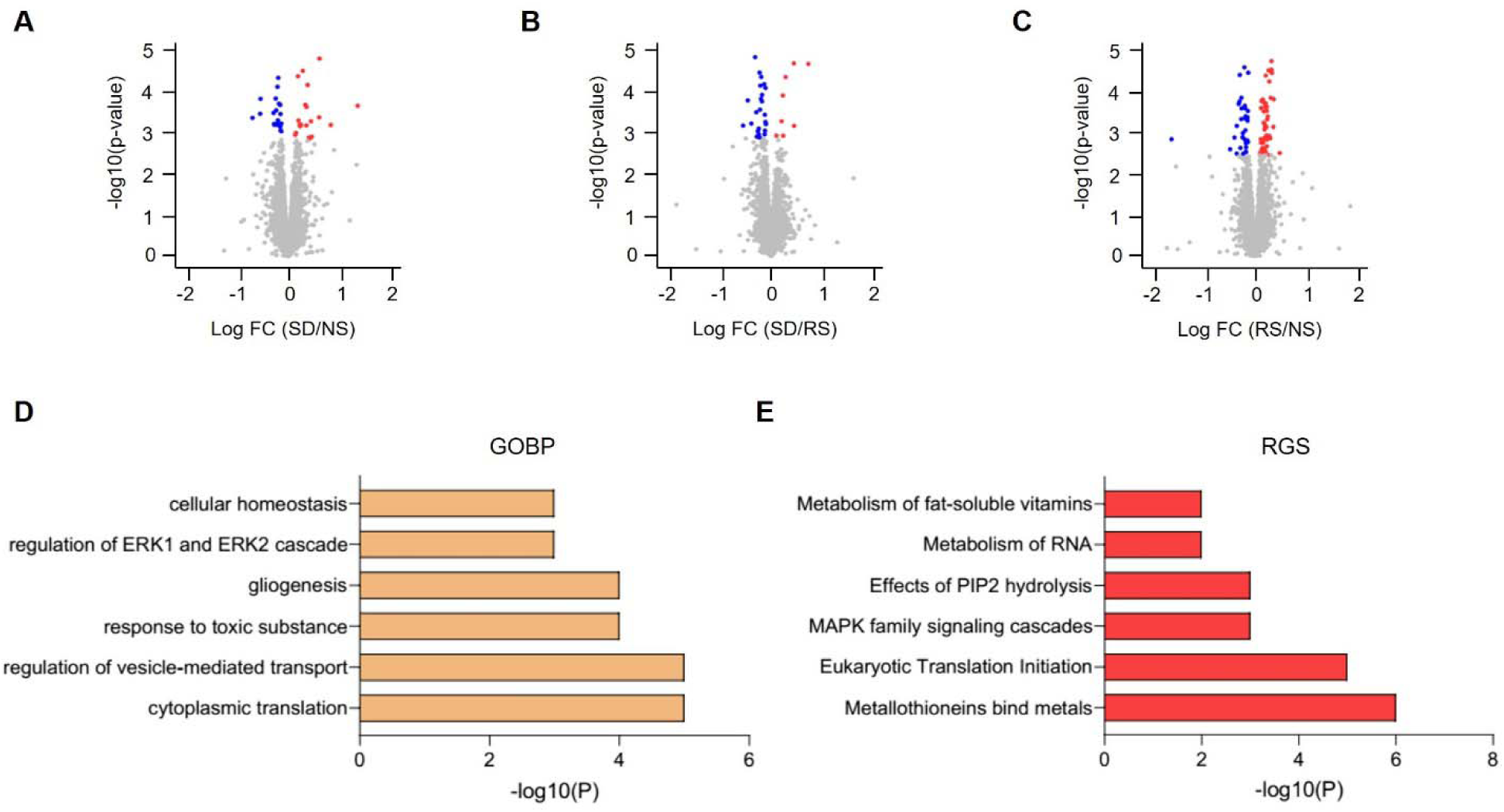
Chronic sleep restriction regulates the global proteome in cerebral cortex. **(A-C)** Volcano plots showing changes in global proteomes of SD/NS (A), SD/RS (B), and (C) RS/NS comparisons. Unpaired t-tests followed by false discovery rate (FDR) analysis were used to compare groups. Significantly upregulated and downregulated protein expression is color-coded with red and blue, respectively (adjusted p-value < 0.2). Proteins in grey are not significantly changed after sleep deprivation. (**D-E**) Bar plots showing significantly enriched top functional annotations from (D) biological processes and (E) reactome. GOBP: gene ontology biological process, RGS: reactome gene sets. NS: normal sleep, SD: sleep deprived, RS: recovery sleep.

To decipher the functional implications of these protein alterations, we conducted gene ontology (GO) analyses of biological processes (GOBP) and reactome gene sets (RGS). GOBP analysis uncovered that CSR modulates key cellular pathways, including cytoplasmic translation, vesicle-mediated transport, metal toxicity responses, gliogenesis, ERK signaling cascades, and cellular homeostasis maintenance (Figure 2D). Through RGS analysis, we identified enrichment in proteins essential for metallothionein binding, eukaryotic translation initiation, MAPK signaling pathways, PIP2 hydrolysis, RNA metabolism, and fat-soluble vitamin processing (Figure 2E). These findings reveal how sleep restriction orchestrates precise molecular changes in cortical function, suggesting a complex interplay between sleep patterns and protein regulation in the brain.

### Aging changes the abundance of cortical proteomes

Next, we explored the molecular and cellular cascades that drive brain aging by conducting comparative proteomic analyses across young (NS), middle-aged (MID), and old (OLD) mice (Figure 3A-C). Our investigation revealed striking alterations in 1,269 unique proteins throughout the aging trajectory (22.1% of 5,737 proteins; q < 0.2). We found that by mid-age (12-14 months), 574 proteins showed significant changes (10% of 5,737; q < 0.2), while this number dramatically increased to 1,132 proteins (19.7% of 5,737; q < 0.2) by 24 months (Figure 3A-B).

**FIGURE 3.**
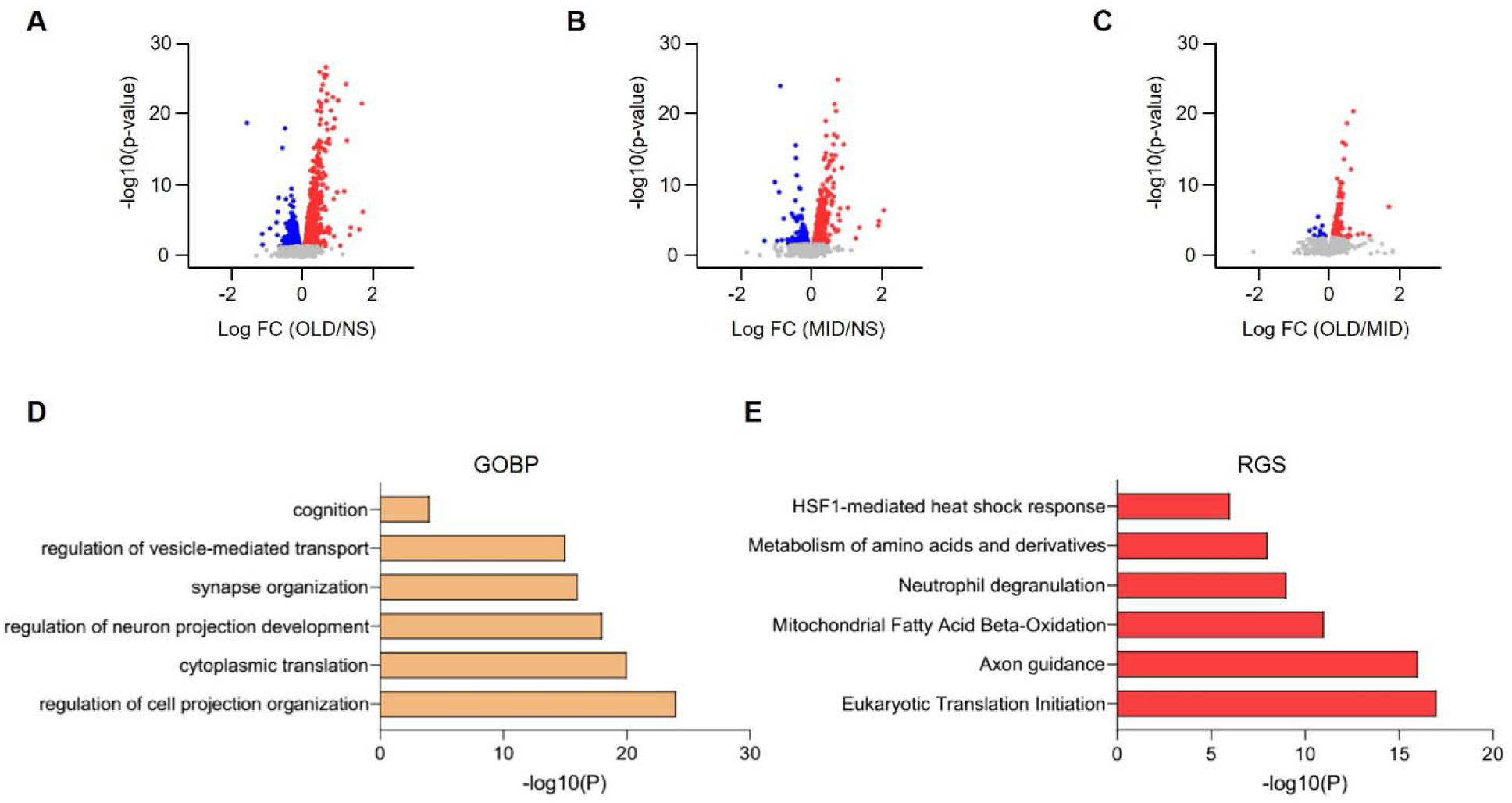
Age-related changes in cortical global proteome. **(A-C)** Volcano plots showing changes in global proteomes of OLD/NS (A), MID/NS (B), and (C) OLD/MID comparisons. Unpaired t-tests followed by false discovery rate (FDR) analysis were used to compare groups. Significantly upregulated and downregulated protein expression is color-coded with red and blue, respectively (adjusted p-value < 0.2). Proteins in grey are not significantly changed after sleep deprivation. (**D-E**) Bar plots showing significantly enriched top functional annotations from (D) biological processes and (E) reactome. GOBP: gene ontology biological process, RGS: reactome gene sets. NS: normal sleep, MID: mid-age, OLD: old age.

To determine the functional significance of these age-dependent protein modifications, we again performed GO and RGS analyses. GOBP analysis unveiled significant age-related changes in critical neural processes, including cell projection organization, cytoplasmic translation, neuron projection development, synapse organization, vesicle-mediated transport regulation, and cognitive function (Figure 3D). RGS demonstrated robust enrichment in proteins governing eukaryotic translation initiation, axon guidance, mitochondrial fatty-acid β-oxidation, neutrophil degranulation, amino acid metabolism, and HSF1-mediated heat shock response (Figure 3E). Together, these results illuminate the molecular landscape of brain aging, revealing systematic alterations in protein networks that orchestrate neural function and plasticity across the lifespan.

### Common proteomic signatures between CSR and aging

To explore potential mechanistic links between chronic sleep restriction (CSR) and aging, we investigated overlapping proteomic signatures across these conditions. Through comparative analysis, we identified 96 proteins exhibiting concordant alterations in both sleep-restricted and aging brains (p < 0.0001) (Figure 4A). We then employed a multi-layered analytical approach – combining gene ontology (GO), pathway analysis, and molecular complex detection (MCODE) –to decode the functional implications of these shared protein modifications.

**FIGURE 4.**
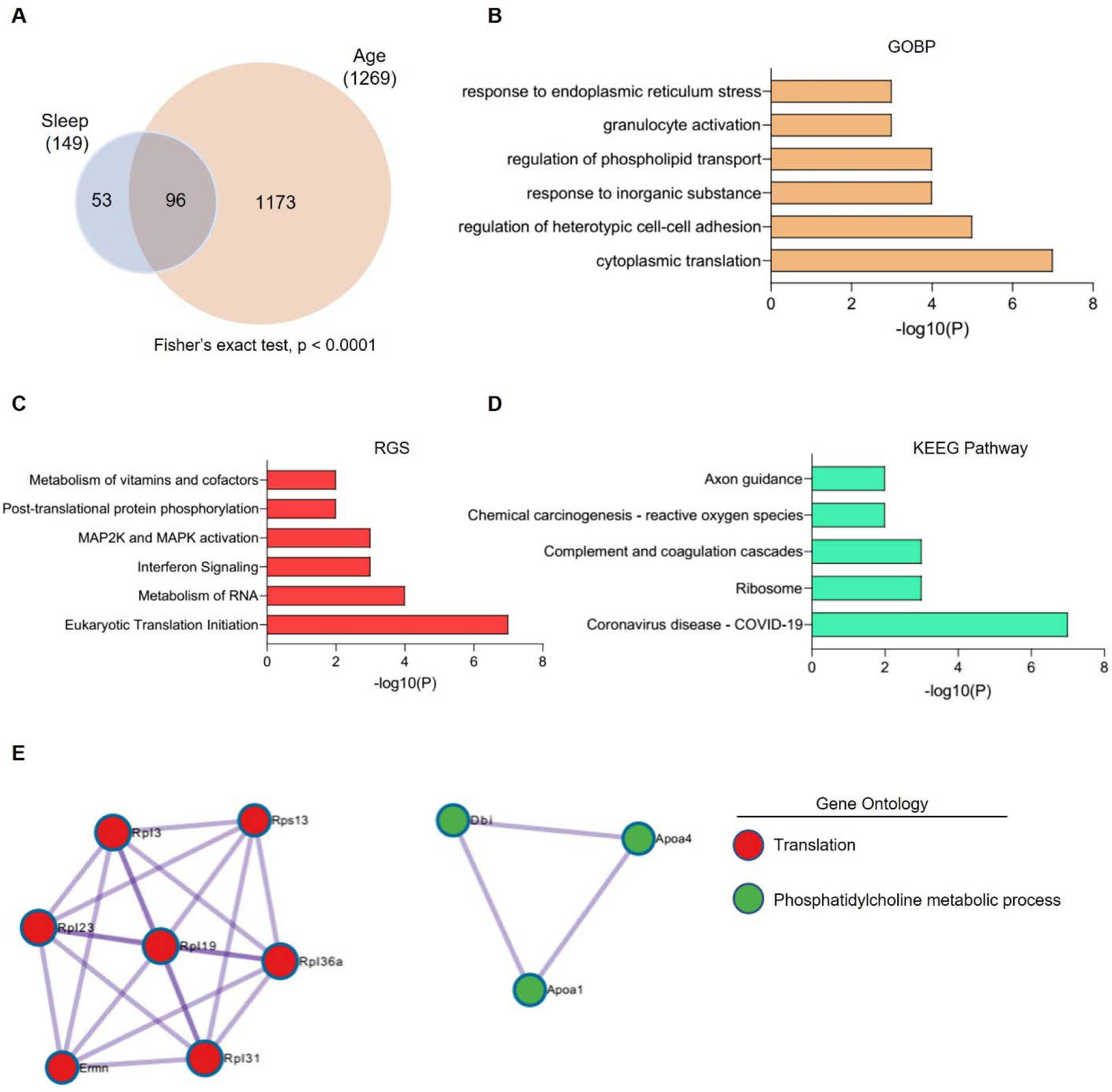
Functional annotations of common proteomic alterations associated with chronic sleep restriction and aging. (**A**) Venn diagrams represent the overlaps between sleep and age-related significant alterations. (**B-D**) Bar plots showing significantly enriched top functional annotations from (B) biological processes, (C) reactome, and (D) KEEG pathways. (**E**) MCODE components (enrichment P-value < 0.001). GOBP: gene ontology biological process, RGS: reactome gene sets, MCODE: Molecular Complex Detection.

GO analysis revealed fascinating enrichment patterns in cytoplasmic translation, heterotypic cell-cell adhesion regulation, inorganic substance responses, phospholipid transport regulation, granulocyte activation, and endoplasmic reticulum stress responses. The reactome analysis further illuminated enrichment in eukaryotic translation initiation, RNA metabolism, interferon signaling, MAP2K and MAPK activation, post-translational protein phosphorylation, and vitamin/cofactor metabolism (Figure 4B-C).

Intriguingly, Kyoto Encyclopedia of Genes and Genomes (KEGG) pathway analysis unveiled significant enrichment in systems linked to coronavirus disease (COVID-19), ribosomal function, complement and coagulation cascades, reactive oxygen species, and axon guidance

(Figure 4D). These findings suggest that both sleep restriction and aging may enhance vulnerability to COVID-19 infection while perturbing complement and coagulation cascades – a mechanism recently implicated in α-synuclein-mediated neurodegeneration and neuronal loss ^20^. We further dissected the complement and coagulation cascade pathways, mapping proteins modified by both CSR and aging (Figure 5). This revealed alterations in key components, including complement component 3 (C3), alpha-2-macroglobulin (A2M), and fibrinogen alpha and beta chains (FGA and FGB). Through protein-protein interaction MCODE analysis, we discovered significant involvement of translation and phosphatidylcholine metabolic processes in both CSR and aging (enrichment p-value < 0.001) (Figure 4E), suggesting shared molecular mechanisms underlying these conditions.

**FIGURE 5.**
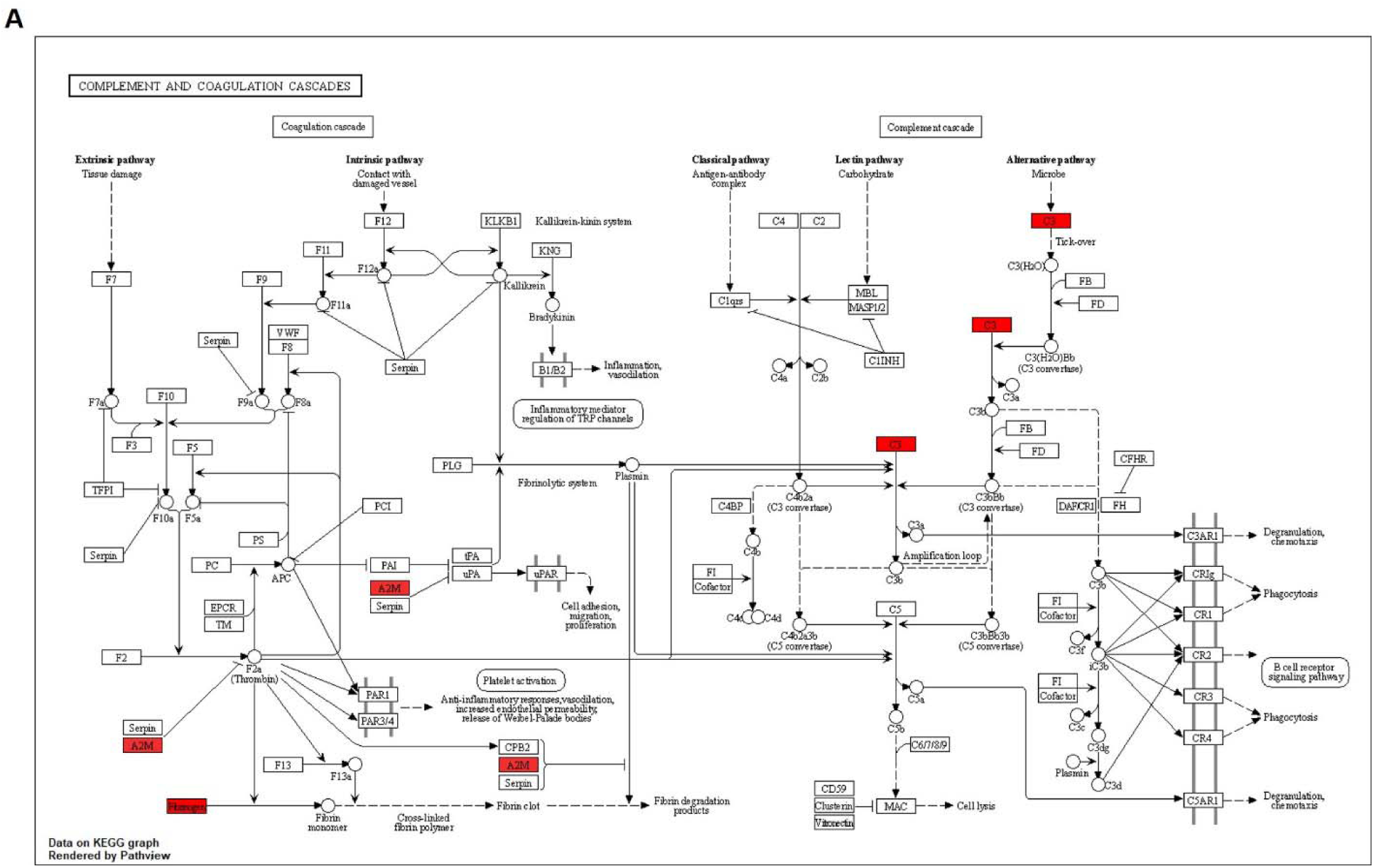
Sleep and age-mediated effects on complement and coagulation cascades. (**A**) KEEG pathway diagram depicting complement and coagulation cascades. Sleep and age-related proteins are colored red.

## DISCUSSION

Mass spectrometry-based proteomics technologies have been widely used to unravel the molecular correlates of certain behaviors and disease conditions ^21–24^. Using this powerful technology, we uncovered crucial molecular links between chronic sleep loss and accelerated brain aging. Our comprehensive analysis identified a significant overlap between proteins altered by chronic sleep restriction (CSR) and aging in the mouse cerebral cortex. The most striking discovery was the shared activation of complement and coagulation cascades in both conditions, suggesting a common molecular pathway through which sleep loss might accelerate brain aging.

In recent years, attempts have been made to elucidate the proteome of sleep-deprived brains, but these studies were mainly aimed at studying the effect of acute sleep deprivation ^18,22,25–27^. In this study, we attempted to mimic real-world conditions of restricted sleep by allowing the mice to sleep only 4 hours for 5 consecutive days and mapped the proteomics alterations in the cerebral cortex. Our differential protein abundance analysis shows that chronic sleep treatments alter only 2.5% of identified proteins. Of note, we recently reported a shift of 22% in neuronal proteome under acute sleep restrictions, although the changes in protein expression in astrocytes were limited to 3% ^18^. These results reveal the variable response of acute and chronic sleep deprivation on protein expression in the brain. Interestingly, this trend is consistent at the level of mRNA expression ^17,19,28,29^. We and others have reported that synapse organization and activity, macromolecule biosynthesis, and energy homeostasis are the prime functional implications of acute sleep disruptions ^17,18,27,30^. On the other hand, sustained sleep loss triggers inflammation and cellular stress ^17^. Here we showed that chronic sleep treatments alter cytoplasmic translation, response to toxic accumulation, MAPK/ERK cascades, gliogenesis, and RNA metabolism. These molecular signatures suggest that CSR particularly impacts non-neuronal cells like astrocytes and microglia, which traditionally serve as support cells in the brain. This finding challenges our traditional neuron-centric view of sleep’s effects on the brain.

Aging is a complex biological process. The speed and trajectory of aging vary from person to person. We furthered our investigation to understand the multifactorial process of brain aging by comparing the proteomics landscape of the cerebral cortex of young, middle, and old-age mice. Age-related proteomic changes were extensive, affecting 19% of identified proteins. These alterations occurred in pathways governing cytoplasmic translation, neuron projection development, synapse organization, vesicular transport, and mitochondrial function. Such widespread changes reflect the complex nature of brain aging, where multiple cellular processes gradually decline over time. Our findings align with previous studies on molecular mechanisms of brain aging ^12,31,32^, confirming that aging affects fundamental cellular processes throughout the brain.

The complement cascade emerged as a key mediator linking sleep loss to accelerated aging. This pathway, part of the innate immune system, normally protects the brain from injury and infection. Both CSR and aging increased the expression of complement component C3, the central molecule in complement activation. C3 drives synapse elimination in the aging central nervous system ^33^, essentially pruning neural connections. Its increased expression appears in age-related disorders like Alzheimer’s and Parkinson’s disease ^20^, suggesting it may contribute to cognitive decline.

Coagulation pathway activation provides another mechanistic link between CSR and aging. We saw the altered expression of fibrinogen chains (FGA, FGB) and alpha-2-macroglobulin (A2M) in both conditions. These proteins normally regulate blood clotting, but in the brain, they can have detrimental effects. Their increased presence contributes to blood-brain barrier disruption and neuroinflammation in aging brains ^34^. A2M expression particularly correlates with early neuronal injury in preclinical Alzheimer’s disease ^35^, marking it as a potential early indicator of brain aging.

The complement and coagulation systems typically maintain brain homeostasis through protective functions, acting to maintain and defend the brain. However, chronic activation of these pathways can trigger neuroinflammation and disrupt synaptic transmission ^36^, much like an overactive immune response causing collateral damage. Our findings suggest that CSR could potentially accelerate brain aging by chronically activating these cascades, essentially wearing down the brain’s protective systems through overuse. This newfound mechanism offers potential therapeutic targets for mitigating the effects of sleep loss and slowing brain aging.

Overall, our study reveals how chronic sleep loss may prematurely age the brain through specific molecular pathways shared between sleep deprivation and aging. Understanding these mechanisms opens new avenues for therapeutic intervention, potentially allowing us to protect the brain from the harmful effects of insufficient sleep. Future research should focus on developing targeted treatments that modulate complement and coagulation cascades to promote healthy brain aging, even in the face of sleep challenges common to modern life.

## MATERIALS AND METHODS

### Animals

All animal studies were carried out in concordance with an approved protocol from the Institutional Animal Care and Use Committee (IACUC) at Perelman School of Medicine at the University of Pennsylvania. Wild-type, male C57BL/6J mice (RRID:IMSR_JAX:000664) were purchased from The Jackson Laboratory and acclimatized in the animal unit for at least 2 weeks before experiments. Mice were selected such that they were aged between 3-4 months (NS, SD, and RS), 12-14 months (MID), and 24 months (OLD) at the time of experiments. Before the chronic sleep restriction protocol began, mice were housed for a week in the sleep cage (Campden/Lafayette Instruments Model 80391) for habituation with ad libitum access to food and water under standard humidity and temperature (21±10°C) on a 12-h light: 12-h dark cycle.

### Sleep deprivation procedures

Sleep deprivation procedures were adapted from the previously described study ^37^. Briefly, sleep deprivation was produced by applying tactile stimulus with a horizontal bar sweeping just above the cage floor (bedding) from one side to the other side of the cage. A five-day procedure of sleep deprivation was performed from ZT4 to ZT20 (20 h) with 4 h of opportunity to sleep and different aged controls were subjected to the same tactile stimulus with a horizontal bar sweeping but twice the speed in their active period from ZT14 to ZT24 (10 h). This ensured that total activity per day was the same across all studied groups. The recovery (RS) group was allowed to recover following SD for two days. Mice had ad libitum access to food and water throughout the experiment. Different aged controls and SD groups were sacrificed on the 6^th^ day and the RS group on the 9^th^ day at ZT4 (Figure 1A). Following sacrifice by cervical dislocation, whole brains were isolated, cortexes quickly dissected, frozen in liquid nitrogen, and stored at −80°C.

### TMT-based quantitative proteomics

The mouse cerebral cortex samples were homogenized in 300μL lysis buffer (50 mM HEPES, 0.5% NP-40, protease, and phosphatase inhibitors) using a pellet pestle (Sigma-Aldrich) for 1 min on ice. Then, mid-sonication was applied for 15 min (30 sec on, 30 sec off; medium power) using a Bioruptor Standard (Diagenode) instrument and lysates were centrifuged at 16,000 × g for 15 min at 4°C. Supernatants were carefully separated and transferred into new microcentrifuge tubes. Protein precipitation was performed with 1:6 volume of pre-chilled (−20°C) acetone overnight at 4°C, followed by centrifugation of lysates at 14,000 x g for 15 minutes at 4°C. Supernatants were discarded without disturbing pellets, and pellets were air-dried for 2-3 minutes to remove residual acetone. Pellets were dissolved in 300 µL 100mM TEAB buffer. Sample processing for TMT-based quantitative proteomics was performed following the same protocol as described previously ^18^. Briefly, each sample’s protein concentration was determined using the Pierce™ BCA Protein Assay Kit (Thermo Fisher Scientific, 23225). 100µg protein per condition was transferred into new microcentrifuge tubes and 5 µL of the 200 mM TCEP was added to reduce the cysteine residues and the samples were then incubated at 55°C for 1-h. Subsequently, the reduced proteins were alkylated with 375 mM iodoacetamide (freshly prepared in 100 mM TEAB) for 30 minutes in the dark at room temperature. Then, trypsin (Trypsin Gold, Mass Spectrometry Grade; Promega, V5280) was added at a 1:40 (trypsin: protein) ratio and samples were incubated at 37°C for 12-h for proteolytic digestion. After in-solution digestion, peptide samples were labeled with 16-plex TMT Isobaric Label Reagents (Thermo Fisher Scientific, A44521) following the manufacturer’s instructions. The reactions were quenched using 5µL of 5% hydroxylamine for 30 minutes. Protein from all five experimental conditions, i.e. NS, SD, RS, MID, and OLD (n=3 biological replicates for each) were labeled with the 15 of the TMT labels within the 16-plex reagent set, while the final mass tag was used for labeling an internal pool containing an equal amount of proteins from each sample. The application of multiplexed TMT reagents allowed the comparison of NS, SD, RS, MID, and OLD samples within the same MS run, eliminating the possibility of run-to-run (or batch) variations.

TMT-labelled samples were resuspended in 5% formic acid and then desalted using a SepPak cartridge according to the manufacturer’s instructions (Waters, Milford, Massachusetts, USA). Eluate from the SepPak cartridge was evaporated to dryness and resuspended in buffer A (20 mM ammonium hydroxide, pH 10) prior to fractionation by high pH reverse-phase (RP) chromatography using an Ultimate 3000 liquid chromatography system (Thermo Scientific). In brief, the sample was loaded onto an XBridge BEH C18 Column (130Å, 3.5 µm, 2.1 mm X 150 mm, Waters, UK) in buffer A and peptides eluted with an increasing gradient of buffer B (20 mM Ammonium Hydroxide in acetonitrile, pH 10) from 0-95% over 60 minutes. The resulting fractions (20 fractions per sample) were evaporated to dryness and resuspended in 1% formic acid prior to analysis by nano-LC MSMS using an Orbitrap Fusion Lumos mass spectrometer (Thermo Scientific).

### Nano-LC Mass Spectrometry

High pH RP fractions were further fractionated using an Ultimate 3000 nano-LC system in line with an Orbitrap Fusion Lumos mass spectrometer (Thermo Scientific). In brief, peptides in 1% (vol/vol) formic acid were injected into an Acclaim PepMap C18 nano-trap column (Thermo Scientific). After washing with 0.5% (vol/vol) acetonitrile 0.1% (vol/vol) formic acid peptides were resolved on a 250 mm × 75 μm Acclaim PepMap C18 reverse phase analytical column (Thermo Scientific) over a 150 minutes organic gradient, using 7 gradient segments (1-6% solvent B over 1 minute, 6-15% B over 58 minutes, 15-32%B over 58 minutes, 32-40%B over 5 minutes, 40-90%B over 1 minute, held at 90%B for 6 minutes and then reduced to 1%B over 1 minute) with a flow rate of 300nL min^−1^. Solvent A was 0.1% formic acid, and Solvent B was aqueous 80% acetonitrile in 0.1% formic acid. Peptides were ionized by nano-electrospray ionization at 2.0kV using a stainless-steel emitter with an internal diameter of 30 μm (Thermo Scientific) and a capillary temperature of 275°C. All spectra were acquired using an Orbitrap Fusion Tribrid mass spectrometer controlled by Xcalibur 4.1 software (Thermo Scientific) and operated in data-dependent acquisition mode using an SPS-MS3 workflow. FTMS1 spectra were collected at a resolution of 120,000, with an automatic gain control (AGC) target of 200,000 and a max injection time of 50 ms. Precursors were filtered with an intensity threshold of 5000, according to charge state (to include charge states 2-7) and with monoisotopic peak determination set to Peptide. Previously interrogated precursors were excluded using a dynamic window (60s +/-10ppm). For FTMS3 analysis, the Orbitrap was operated at 50,000 resolutions with an AGC target of 50,000 and a max injection time of 105 ms. Precursors were fragmented by high energy collision dissociation (HCD) at a normalized collision energy of 60% to ensure maximal TMT reporter ion yield. Synchronous Precursor Selection (SPS) was enabled to include up to 5 MS2 fragment ions in the FTMS3 scan.

### Database search and statistical analysis of quantitative proteomics data

Quantitative proteomics raw data files were analyzed using the MaxQuant (version 2.1.3.0) ^38^. MS2/MS3 spectra were searched against UniProt database specifying Mus musculus (Mouse) taxonomy (Proteome ID: UP000000589; Organism ID: 10090). All searches were performed using “Reporter ion MS3” with “16-plex TMT” as isobaric labels with a static modification for cysteine alkylation (carbamidomethylation), and oxidation of methionine (M) and protein N-terminal acetylation as the variable modifications. Trypsin digestion with a maximum of two missed cleavages, minimum peptide length of seven amino acids, precursor ion mass tolerance of 5 ppm, and fragment mass tolerance of 0.02 Da were specified in all analyses. The false discovery rate (FDR) was specified at 0.01 for peptide spectrum match (PSM), protein, and site decoy fraction. TMT signals were corrected for isotope impurities based on the manufacturer’s instructions. Subsequent processing and statistical analysis of quantitative proteomics datasets were performed using Perseus (version 2.0.6.0) ^39^ and DEseq2 ^40^. During data processing, reverse and contaminant database hits and candidates identified only by site were filtered out. Finally, unpaired t-tests followed by multiple-testing adjustment (Benjamini-Hochberg method) were performed to compare sleep and aged control groups.

### Gene ontology (GO) analysis

Significantly and differentially abundant proteins across sleep treatments and different age groups were subjected to GO analysis ^41^. Significantly enriched GO terms were visualized as bar plots for GOBP and RGS. Protein-protein interactions network analysis was performed with MCODE. We visualized the enriched KEGG pathway by using clusterProfiler and pathview^42,43^.

### Statistical methods

All experimental subjects are biological replicates. R or GraphPad Prism 8 software was used to perform statistical tests and plots. Unpaired t-tests were performed to compare sleep treatments and aged control groups. For this analysis, we used multiple-testing adjusted (Benjamini-Hochberg) q-values < 0.2, unless indicated otherwise. Detailed sample sizes, statistical tests, and results are reported in the figure legends.

## Acknowledgments

A.B.R. acknowledges funding from the Perelman School of Medicine, the University of Pennsylvania, and the Institute for Translational Medicine and Therapeutics (ITMAT) at the University of Pennsylvania. This work was supported also by NIH DP1DK126167 and R01GM139211 (A.B.R.).

## Author contributions

P.K.J. and A.B.R. conceived and designed the experiments. P.K.J. performed mouse sleep deprivation experiments, tissue dissections, and sample processing. P.K.J. and U.K.V. analyzed proteomics data. A.B.R. supervised the entire study and secured funding. The manuscript was written by P.K.J. and A.B.R. with input from U.K.V. All authors agreed on the interpretation of data and approved the final version of the manuscript.

## Declaration of Interests

The authors declare no competing interests.

## Funding Declaration

This work was supported also by NIH DP1DK126167 and R01GM139211 (Akhilesh B. Reddy).

## Notes

### Competing Interest Statement

The authors have declared no competing interest.

